# Developmental emergence of first- and higher-order thalamic neuron molecular identities

**DOI:** 10.1101/2024.01.22.576610

**Authors:** Quentin Lo Giudice, Robin J. Wagener, Philipp Abe, Laura Frangeul, Denis Jabaudon

## Abstract

The thalamus is organized into nuclei that have distinct input and output connectivities with the cortex. While first-order (FO) nuclei – also called core nuclei – relay input from sensory organs on the body surface and project to primary cortical sensory areas, higher-order (HO) nuclei – matrix nuclei – instead receive their driver input from the cortex and project to secondary and associative areas within cortico-thalamo-cortical loops. Input-dependent processes have been shown to play a critical role in the emergence of FO thalamic neuron identity from a ground state HO neuron identity, yet how this identity emerges during development remains unknown. Here, using single-cell RNA sequencing of the developing embryonic thalamus, we show that FO thalamic identity emerges after HO identity and that peripheral input is critical for the maturation of excitatory, but not inhibitory FO-type neurons. Our findings reveal that subsets of HO neurons are developmentally co-opted into FO-type neurons, providing a mechanistic framework for the diversification of thalamic neuron types during development and evolution.

**Summary Statement:** Subsets of higher-order thalamic neurons are developmentally co-opted into first-order type neurons, providing a framework for the diversification of thalamic neuron types during development and evolution.

## Introduction

The thalamus is most often referred to in relation to its function as a relay from sensory organs in the periphery *–* the retina, cochlea, skin mechanoreceptors *–* to their corresponding target area in the neocortex *–* the primary visual, auditory, and somatosensory cortices *–* (Clascá et al., 2012; López-Bendito and Molnár, 2003). Such relay function is mediated by so-called first-order (FO) thalamic nuclei, namely the dorsolateral geniculate nucleus (LG) for vision, the ventrobasal nucleus (VB) for somatosensation, and the ventral medial geniculate nucleus (vMG) for audition.

Most of the thalamus, however, does not serve such functions, but instead acts within cortico-thalamo-cortical loops. In these loops, input from layer 5b neurons in primary sensory areas is forwarded to other cortical areas (secondary sensory areas, associative areas), allowing integration of sensory information and sensorimotor modulation (Antón-Bolaños et al., 2018; Jabaudon, 2017; Theyel et al., 2009). As examples, the higher-order (HO) nucleus for vision is the latero-posterior nucleus (LP), while the posteromedial nucleus (Po) is the HO nucleus for somatosensation. Hence, for each sensory modality, there exists a cortico– thalamo–cortical loop in which primary sensory cortex receives input from exteroceptive FO nucleus, and projects back to the corresponding HO thalamic nucleus, itself targeting a secondary sensory cortex (Fig. 1A).

**Fig. 1.**
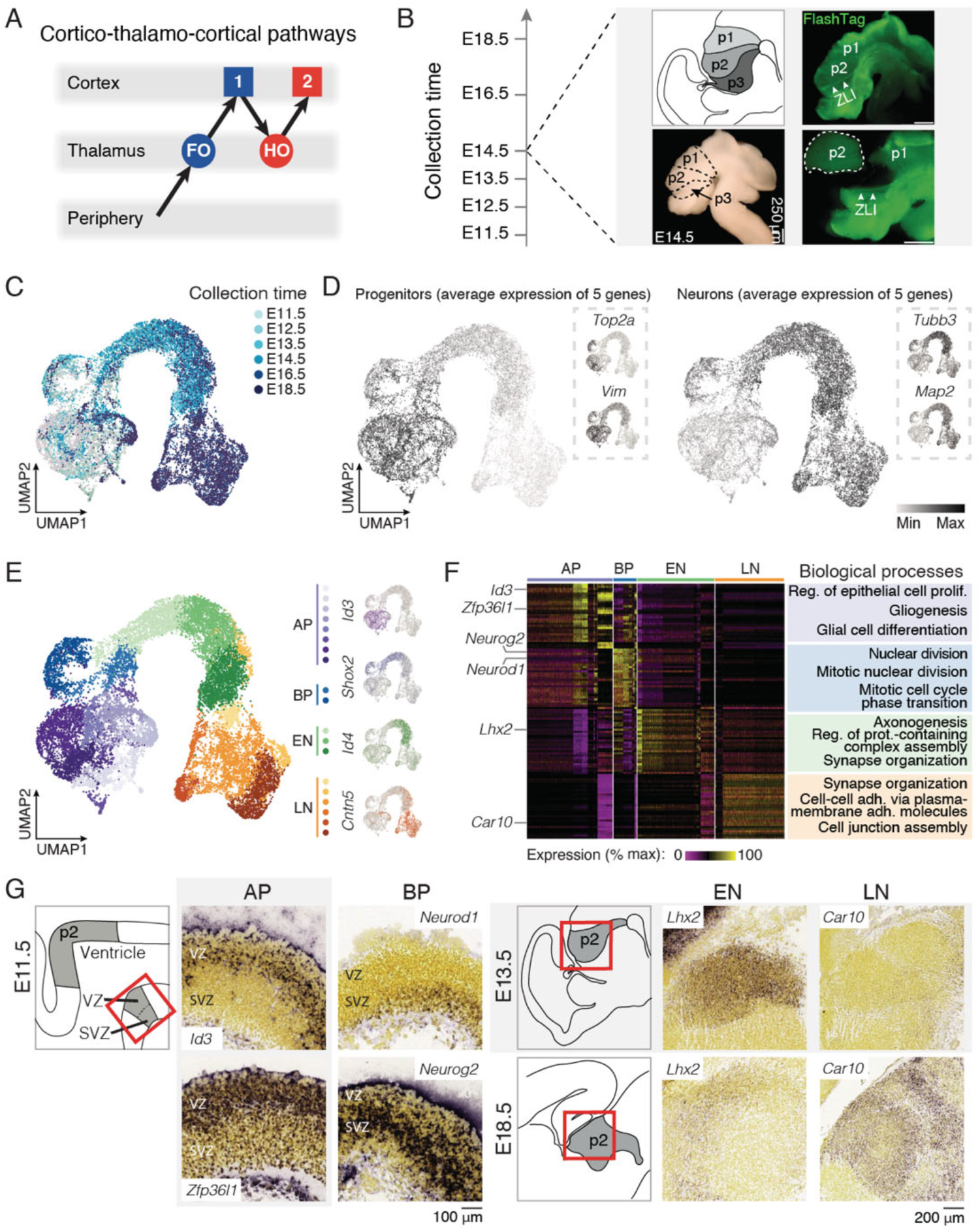
Thalamic stage-dependent transcriptional diversity. (A) Schematic representation of cortico-thalamo-cortical pathways. (B) Schematic illustration of the experimental timeline and the procedure for microdissection. (C) UMAP representation of the single-cell RNA sequencing dataset color-coded by age of collection. (D) Progenitors and neurons can be distinguished by their combinatorial expression of key marker genes (*n* = 5 transcripts with *Btg2*, *Pax6*, *Vim*, *Hes1* and *Nes* for progenitors, and *Map2*, *Tubb3*, *Eno2*, *Rbfox3* and *Nefl* for neurons). (E) Cluster analysis reveals transcriptionally distinct and temporally dynamic cellular clusters (left): apical progenitors (AP) cells in purple gradient, basal progenitors (BP) cells in blue gradient, early neurons (EN) cells in green gradient and late neurons (LN) cells in orange gradient. Individual cell type representative genes are shown in their respective color (right). (F) Expression of the top 30 most expressed genes by cellular cluster highlight cellular diversity (left). Example of biological processes of gene ontologies associated with each cellular cluster. (G) Schematic representation of the p2 domain at E11.5, E13.5 and E18.5 and *in situ* hybridization (ISH) sections of selected differentially expressed genes showing distinctive expression in AP, BP, EN and LN; image source: Allen Developing Mouse Brain Atlas (developingmouse.brain-map.org). FO, first order; HO, higher order; SVZ, subventricular zone; VZ, ventricular zone; ZLI, zona limitans intrathalamica. Scale bars: 250 μm (A); 100 μm (G, left); 200 μm (G, right).

Recent findings have identified the molecular identities of developing thalamic neurons and shed light on their clonal relationships, yet how this relates to the distinction between FO and HO neuron identity remains unclear (Gezelius et al., 2017; Govek et al., 2022; Moreno-Juan et al., 2017; Shi et al., 2017; Yuge et al., 2011). Input-dependent processes appear to play a role in this process since input ablation (e.g. vibrissectomy or enucleation) performed at critical periods of early postnatal development dramatically affects the development of FO thalamic nuclei and their corresponding primary cortical target (Frangeul et al., 2016; Giasafaki et al., 2022; Grant et al., 2016; Van der Loos and Woolsey, 1973). This is visible in terms of morphological organization, whereby infraorbital nerve section disrupts the somatotopic map of the whiskers in the VB (Frangeul et al., 2014; Frangeul et al., 2016; Jensen and Killackey, 1987; Van der Loos and Woolsey, 1973) or enucleation disrupts retinotopic map and interneuron migration in the LG (Golding et al., 2014; Wiesel and Hubel, 1963). Such effects, however, are less visible in HO nuclei, consistent with their circuit position, which is more remote from input from the periphery (Frangeul et al., 2014; Frangeul et al., 2016; Theyel et al., 2009).

In molecular terms, input ablation at early postnatal ages in mice leads to an HO nucleus identity in FO nuclei, both in terms of molecular identity and connectivity (Frangeul et al., 2016; Grant et al., 2016). Likewise, genes normally enriched in LG during development are preferentially downregulated following monocular enucleation (Giasafaki et al., 2022). While these results support the idea that HO-type genetic identity is a developmental ground-state feature of thalamic neurons and the FO identity is acquired secondarily in an input-dependent manner, this has not been directly tested and is the topic of the present study.

Here, using single-cell and single-nucleus RNA sequencing of the developing thalamus between E11.5 and E18.5, we first identify the timeline of the emergence of FO and HO neuron molecular identity and, within these classes, that of LG, LP, VB, and Po identities, and identify transcriptional programs that are conserved and distinct between these cells. We show that FO identity emerges after HO identity, and that enucleation prevents the normal maturation of LG neurons in a cell-type specific manner, while LP neurons are only minimally affected. Our results establish a mechanistic framework for the diversification of thalamic neuron types during development, in which FO nuclei emerge from a specialization of HO nuclei, with peripheral input playing a cell-type specific role in this process.

## Results

### Identification of the developmental transcriptional programs of thalamic nuclei

Essentially all thalamic neurons are born between embryonic day (E) 11.5 and E14.5 (see https://neurobirth.org/ and Fig. S1) (Baumann et al., 2023). As a first foray into the developmental molecular diversity of FO and HO nuclei, we performed single-cell RNA sequencing of prosomere 2 (p2), from which the thalamus arises, at E11.5, E12.5, E13.5, E14.5, E16.5, and E18.5. In select experiments, we performed FlashTag labeling (Govindan et al., 2018) to label structure boundaries – in particular, the zona limitans intrathalamica – allowing precise microdissection (Fig. 1B). Following tissue collection and dissociation, 10X single-cell RNA-sequencing, quality control and data filtering (see Methods), we obtained a total of 18’977 transcriptionally characterized cells, which we displayed in 2 dimensions using UMAP dimensionality reduction (Fig. 1C-E). This approach revealed that cells were molecularly organized based on their type and differentiation state. Progenitors were grouped together and were the predominant cell type until E13.5, when neurogenesis is largely complete, while neurons progressively emerged throughout time, reflecting progressive maturation of the age-specific cohorts (Fig. 1C). This developmental timeline was confirmed using markers of progenitors and postmitotic neurons, respectively (Fig. 1D, left, and right). Unbiased cluster analysis revealed a total of 19 clusters that could be functionally grouped into 4 classes of cells (Fig. 1E): apical progenitors, expressing *Nes* and *Sox2*, basal progenitors, which expressed *Neurog2* and *Btg2* (Moreau et al., 2021; Telley et al., 2016), and early and late neurons, which expressed classical markers of neuronal differentiation *Tubb3* and *Id4* (Menezes and Luskin, 1994; Yun et al., 2004) and maturity *Eno2* and *NeuN* (Duan et al., 2016; Schmechel et al., 1978), respectively (Table S1, Fig. S2). Interestingly, two types of cells were found to be cycling, i.e. apical progenitors and basal progenitors (Table S2, Fig. S3) (Wang et al., 2011). Analysis of differentially expressed genes across clusters and cell types unveiled the sequential functional transcriptional programs at play in the developing thalamus: initially, genes involved in regulation of cell division and proliferation predominated with transcripts such as *Cdca8* and *Cenpf* respectively, while axon and synapses-related genes were expressed in early and late neurons with transcripts such as *Epha3* and *Dlg2* respectively (Fig. 1F, Table S1, Table S3). Finally, the specificity of the genes identified was confirmed using the Allen Developing Mouse Brain Atlas *in situ* database (developingmouse.brain-map.org). AP- and BP-specific genes were enriched in the germinal zones of the developing thalamus (in the ventricular and subventricular zones, respectively). EN- and LN-specific genes were expressed in the mantle zone of prosomere 2, at early and late developmental stages, respectively (Fig. 1G, Fig. S2). Together, these data identify the core molecular cell types present in the developing thalamus.

To get a finer-grained understanding of the sequential molecular programs that are unfolding in thalamic cells during this developmental period, we used a pseudo-differentiation approach (here called “pseudotime”, for simplicity). Across this axis, we identified transcriptional waves of sequential gene expression, i.e. genes which were most dynamically expressed across time (Fig. 2A). This approach revealed four main waves of transcription for a total of 268 genes, that progressed in function from control of cell cycle properties, followed by axonogenesis, cellular maturation and microtubule organization (Fig. 2B,C Table S4 and Table S5). Here again, validation using available in situ datasets for select genes confirmed the temporal progression observed with the single-cell analysis (Fig. 2D). Together, these data identify the temporal transcriptional programs at play in the developing thalamus.

**Fig. 2.**
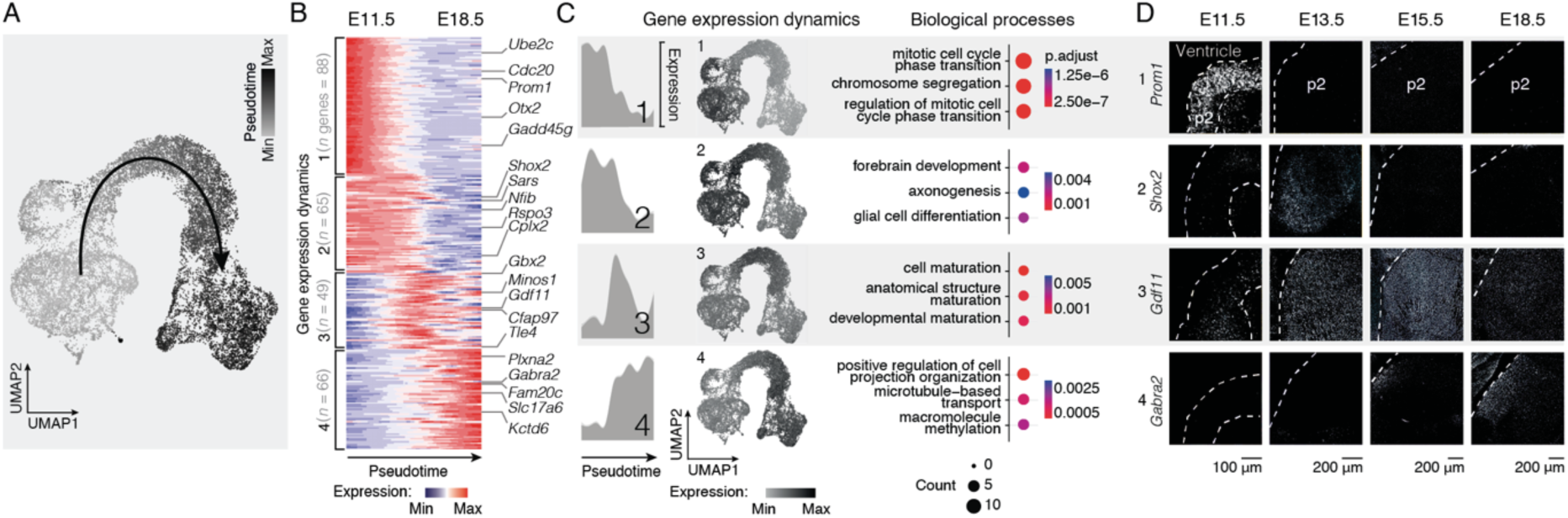
Core thalamic developmental programs are sequentially expressed in four molecular waves. (A) Pseudotime values on the UMAP embedding showing the developmental continuum-like organization of cells. (B) Unbiased clustering of genes based on expression dynamics reveals four distinct transcriptional waves sequentially expressed along the pseudotime axis. Example of selected differentially expressed genes are provided for each wave (left). (C) The four distinct transcriptional waves reveal sequential expression peaks (left). Average of expression of each wave shows discrete pattern of expression along the UMAP representation following the pseudotime axis (middle). Example of biological processes of gene ontologies associated with each transcriptional wave (right). (D) In situ hybridization (ISH) sections of selected differentially expressed genes showing distinctive expression in the different waves; image source: Allen Developing Mouse Brain Atlas (developingmouse.brain-map.org). Scale bars: 100 μm (D, E11.5); 200 μm (D, E13.5, E15.5, E18.5).

### Emergence of FO and HO neuron identity

We next examined how FO and HO molecular identity emerged during development. Previous work has identified several gene markers that distinguish these two types of nuclei in the early postnatal brain and adulthood (Frangeul et al., 2016) (Table S6), yet when and how these identities emerge remains unclear. A limiting factor to bioinformatically fate-map FO- and HO-type neurons based on adult markers is the fact that they are not necessarily expressed at earlier stages of development. To circumvent this limitation, we designed an approach in which we ranked cells based on their relative expression of classical mature FO and HO marker genes and defined the top-ranked cells as “seed cells” from which we identified new differentially expressed genes to serve as markers for analysis in earlier developmental ages (Fig. 3A). Cells were defined as FO- or HO-type based on above-threshold expression of the new gene set (Fig. 3B and Table S7). This analysis revealed that HO identity emerges earlier and over a more prolonged period than that of FO identity, the latter being restricted to a transient period at the end of the time window (Fig. 3C). We then identified differentially expressed genes in FO and HO-type cells during development. This revealed a prolonged early wave of gene expression in HO neurons, consisting of around 1’350 early and 700 late genes, while FO neurons instead progressed rapidly to more mature states with around 550 early and 1’300 late genes (Fig. 3D, Fig. S4 and Table S8). When genes expressed both by FO and HO were examined, their time course was more transient in FO than HO neurons, suggesting different paces of maturation (Fig. S4C). Genes expressed specifically by FO neurons included *Nefm*, *Actb* and *Rps29*, which are involved in neurofilament organization, axonogenesis, and ribosomal biogenesis respectively. Genes expressed specifically by HO-type cells included *Robo2*, *Alcam* and *Grin2a*, which are involved in biological processes such as axon guidance, cell adhesion, and synaptic transmission respectively (FO, n = 861 genes; HO, n = 723 genes; Fig. S5, Table S7 and Table S9). The specificity of the genes identified was confirmed using an available spatial transcriptomic datasets (Zhang et al., 2023): FO-specific genes were enriched in FO nuclei of the thalamus, while HO-specific genes were mostly expressed in HO nuclei of the thalamus (Fig. 3E). Together, these data reveal that HO molecular identity emerges mostly prior to FO molecular identity and identify the transcriptional programs at play during the development of these two types of thalamic nuclei.

**Fig. 3.**
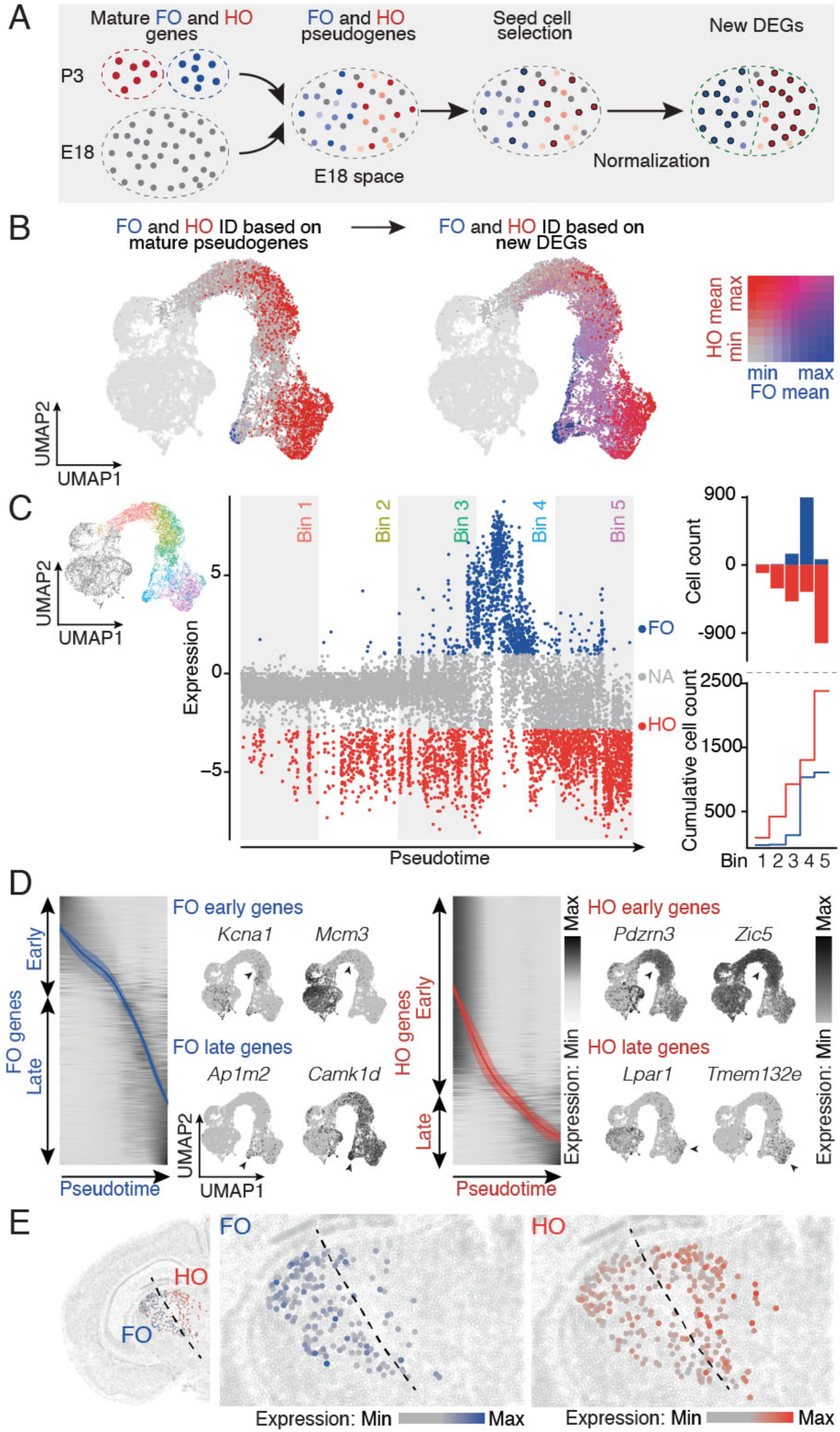
Emergence of thalamic first and high order identities occurs at distinct paces. (A) Schematic illustration of the strategy used for first order (FO) and higher order (HO) final cells selection during embryonic development. (B) UMAP representation of FO and HO mature pseudogenes (left, 15 FO genes, 15 HO genes from previously published data Frangeul et al., 2016, see Table S5) allowing to predict FO and HO differentially expressed genes during embryonic development (right). (C) Each post-mitotic cell is assigned with its FO/HO ratio score (left). Distribution of the cells into 5 consecutives bins and cumulative plot show an early emergence of the FO identity compared to HO (right). (D) Specific transcriptional waves for FO and HO lineages and example feature plot of select genes are provided for each wave. (E) Spatial transcriptomic of select differentially expressed genes (spatial transcriptomic dataset from Zhang et al., 2023). FO, first order; HO, higher order.

### Emergence of VB, LG, Po, and LP neuron identity

Building on the strategy above, we next examined the emergence of modality-specific FO and HO nuclei, namely the VB and Po for the somatosensory system and the LG and LP for the visual system (Fig. 4). Using the seed-cell-approach, but this time with VB- and Po-specific genes (Fig. 4A), and LG- and LP-specific genes (Fig. 4B, Table S10 and Table S11), we examined the time course of emergence of neurons with these identities (Fig. 4C,D). Consistent with temporally shifted differentiation programs, the unfolding of the gene expression programs associated with VB or LG neuron identity, on the one hand, and Po and LP neuron identity, on the other hand, were shifted in time, with HO-nuclei programs unfolding more rapidly, particularly in LP vs. LG (Fig. 4E,F and Table S12). Based on the analyses above, the final identity of these four nuclei could be established, allowing definition of nucleus-specific markers (Fig 4G and Table S11). Common ontologies shared amongst the four nuclei included synaptic assembly (Dscam, Nrg3, Pclo), neurotransmitter secretion (Cacna1b, Nrxn1, Rims1) and nervous system development (Gabrb3, Dclk1, Nrg3) (Fig. S6, Table S11 and Table S13). An available spatial transcriptomic dataset (Zhang et al., 2023) revealed that these newly identified genes had a spatial distribution appropriate for their identity (Fig. 4H). Hierarchical clustering of cells with VB, Po, LG, LP identities revealed stronger similarities within hierarchical levels (i.e. VB and LG vs. Po and LP, Fig. 4I), than within modalities (i.e VB and Po vs. LG and LP) consistent with previous results in the mature thalamus (Frangeul et al., 2016). Together, these data reveal that LP molecular identity emerges prior to LG molecular identity, and, to a lesser extent, Po prior to VB, by identifying new molecular makers for these four thalamic nuclei during development.

**Fig. 4.**
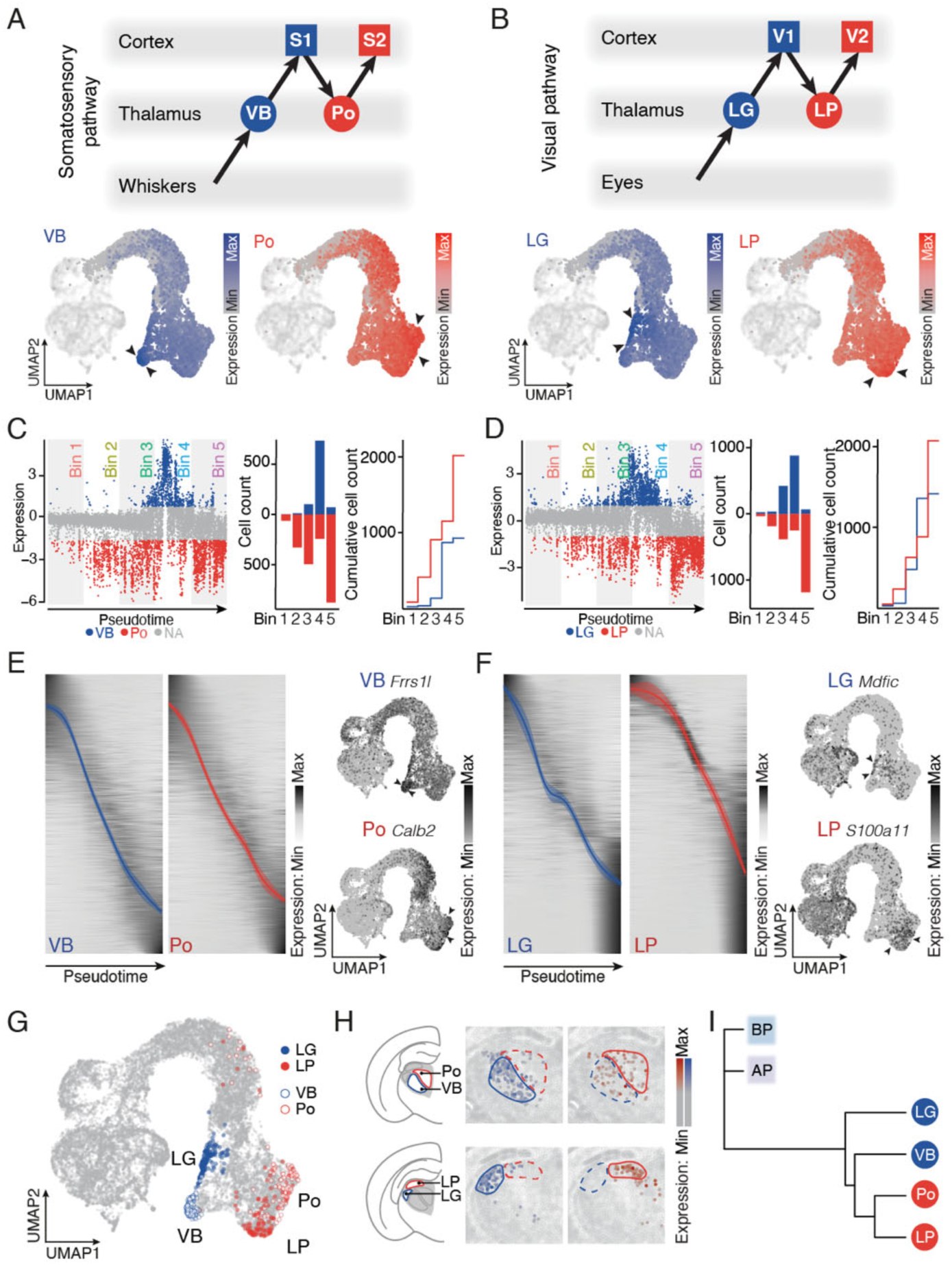
Acquisition of nuclei-specific identities during development relies on both shared and distinct molecular programs. (A) Schematic representation of somatosensory pathway (top). UMAP representation of VB and Po differentially expressed genes during embryonic development using the same strategy as in Fig. 3B (bottom, 10 VB genes, 10 Po genes from previously published data (Frangeul et al., 2016, see Table S10). (B) Schematic representation of visual pathway (top). UMAP representation of LG and LP differentially expressed genes during embryonic development using the same strategy as in Fig. 3B (bottom, 10 LG genes, 10 LP genes from previously published data Frangeul et al., 2016, see Table S10). (C) Each post-mitotic cell is assigned with its VB/Po ratio score (top). Distribution of the cells into 5 consecutive bins and cumulative plot show early emergence of VB identity compared to Po (bottom). (D) Each post-mitotic cell is assigned with its LG/LP ratio score (top). Distribution of the cells into 5 consecutive bins and cumulative plot show an early emergence of the LG identity compared to LP (bottom). (E) Specific transcriptional waves for VB and Po lineages and example feature plot of select genes are provided for each wave. (F) Specific transcriptional waves for LG and LP lineages and example feature plot of select genes are provided for each wave. (G) UMAP representation of VB, Po, LG and LP cells for late timepoints after selection process. (H) Spatial transcriptomic of select differentially expressed genes showing distinctive expression in VB, Po, LG and LP thalamic nuclei (spatial transcriptomic dataset from Zhang et al., 2023). (I) Hierarchical clustering depicting the LG/LP/VB/Po organization within the framework of AP/BP/EN/LN cell categories. LG, dorsolateral geniculate nucleus; LP, pulvinar/latero-posterior nucleus; Po, posteromedial nucleus; S1, primary somatosensory cortex; S2, secondary somatosensory cortex; V1, primary visual cortex; V2, secondary visual cortex; VB, ventrobasal nucleus.

### Input-dependent differentiation of the LG is neuron-type-specific

Focusing on visual pathways, we examined the extent to which the developmental transcriptional programs of the LP and LG were driven by input-dependent processes after birth (Fig. 5). For this purpose, we performed bilateral enucleation at P0, collected LG and LP at P3 and P7, and using single-cell RNA sequencing, assessed and compared molecular identities across ages and conditions (Fig. 5A).

**Fig. 5.**
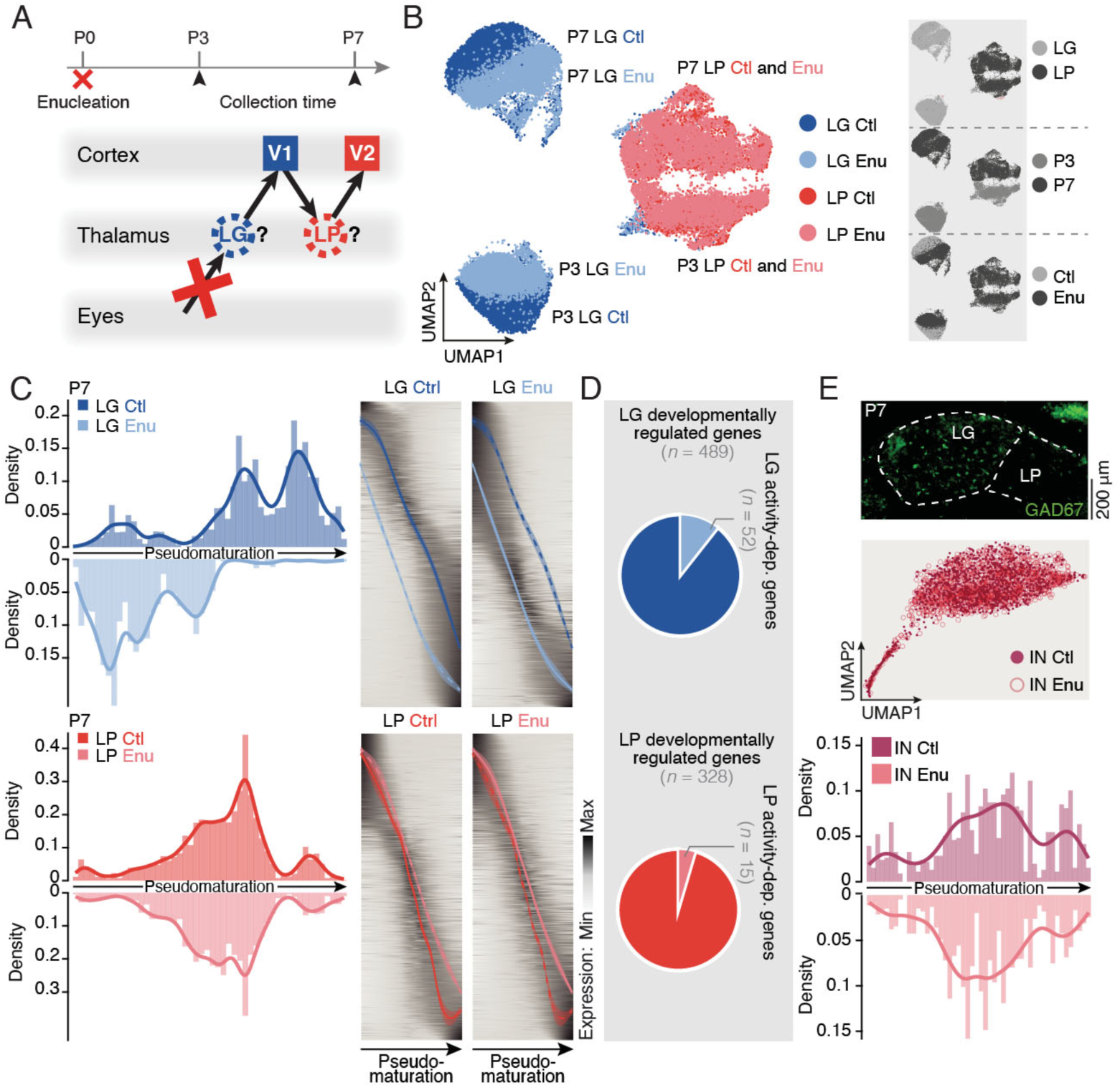
Cell-type-specific transcriptional responses to input deprivation. (A) Schematic illustration of the experimental timeline and the procedure for microdissection. (B) UMAP representation of the single-cell RNA sequencing dataset color-coded by condition (LG Ctl, dark blue; LG Enu, light blue; LP Ctl, dark red; LP Enu, light red). Insets are UMAP representation of the different condition LG/LP (top right), P3/P7 (middle right) and Ctl/Enu (bottom right). (C) Each LG Ctl and LG Enu (top left), and LP Ctl and LP Enu (bottom left) cell is placed along the pseudotime axis. Specific transcriptional waves for LG Ctl and LG Enu (top right), and LP Ctl and LP Enu (bottom right) lineages are represented. (D) Pie chart representing the number of LG (top, *n* = 52) or LP (bottom, *n* = 15) activity-dependent genes among developmentally regulated genes (LG, *n* = 489; LP, *n* = 328). (E) Coronal section in GAD67^GFP^ mice showing interneurons (Ins) in the LG (top). UMAP representation of the IN in Ctl and Enu conditions (middle). Each IN Ctl and IN Enu cell is placed along the pseudotime axis (bottom). Ctl, control; Enu, Enucleation; IN, interneuron; LG, dorsolateral geniculate nucleus; LP, pulvinar/latero-posterior nucleus; V1, primary visual cortex; V2, secondary visual cortex. Scale bar: 200 μm (E).

Focusing first on glutamatergic neurons (Fig. 5B), we found that while there was a striking shift in the identities of LG neurons at these two ages (light and dark blue cell clusters in the UMAP), LP neuron identity remained largely unaffected (light and dark red cells in the UMAP), confirming and extending previous results Frangeul et al., 2016).

Analysis of differentially-expressed genes revealed a downregulation of *Dcc*, a netrin receptor, in the LG after enucleation, while another axon guidance molecule, *Robo2*, was upregulated. Interestingly, Dcc/Netrin signaling and Robo/Slit signaling have distinct roles in axonal guidance, which could provide a basis for the distinct target specificities of FO and HO axons (Deiner et al., 1997; Lo Giudice et al., 2019; Thompson et al., 2009; Zhang et al., 2012). Downregulated genes after enucleation were involved in cholinergic synapse and calcium signaling while ontology upregulated genes GABAergic synapse and glutamatergic synapse. In LP neurons, *Lrrc4c*, a netrin G1 ligand, was downregulated after enucleation, while *Il1rapl2*, a member of the interleukin 1 receptor family playing a role at synapses, was upregulated. Due to the low number of differentially expressed genes in the LP between conditions, the only relevant ontologies were modulation of synaptic transmission and homo/heterophilic cell adhesion via plasma membrane adhesion molecules (Fig. S7, Fig. S9, Table S14 and Table S15).

We next examined how input deprivation affects the temporal unfolding of transcriptional programs in LG and LP neurons using the pseudotime alignment approach previously applied. This revealed that input-deprived LG neurons lagged in their differentiation compared to control LG neurons, highlighting a critical role for retino-thalamic input in driving the maturation of glutamatergic neurons in this nucleus (Fig. 5C). In the LG, about ten percent (10.6%) of developmentally regulated genes were affected by input deprivation, while this was the case for less than five percent (4.6%) of developmentally regulated LP genes (Fig. 5D). Within the LG, different subclusters of neurons were identified, with different sensitivities to enucleation, perhaps relating to the precise nature of the input these cells were receiving (Fig. S9, Table S16). Finally, in contrast to the input-dependent sensitivity of glutamatergic neuron differentiation, when inhibitory interneurons were examined, the temporal unfolding of their transcriptional programs was unaffected, revealing that even within a single nucleus, cell-type specific, input-dependent processes are at play (Fig. 5E).

## Discussion

Our findings reveal that HO neuron identity is a developmental ground state, while FO neuron identity emerges later. Upon input deprivation in the visual system, the differentiation of FO glutamatergic neurons is delayed, while HO identity is not detectably affected. Similarly, inhibitory neurons are not detectably affected. Thus, even within single thalamic nuclei, neuronal types have distinct susceptibilities to input-dependent differentiation.

A fundamental difference between FO and HO inputs is the topographical organization of their synaptic inputs. Hence, while the PrV nucleus of the trigeminal nerve projects in a somatotopic manner to the VB, projections to the PO are, instead, less precise, and a corresponding difference exists for projections of VB and PO nuclei to their cortical targets (Herkenham, 1980; Pouchelon et al., 2012). Similarly, HO-type projections in the cortex appear to rely mostly on metabotropic synaptic transmission, while ionotropic channels mediate FO transmission, arguably allowing for a more precise temporal coding of information. These observations may account for the fact that FO nuclei (here, the LG) are more sensitive than HO nuclei (here, the LP) to input deprivation.

The observation that interneurons are not detectably transcriptionally affected by input deprivation is novel and may reflect the fact that this cell type invades the LG relatively late, during the first few postnatal days. We have, however, previously shown that enucleation affects the migrations of these cells within this nucleus, as well as their synaptic integration within visual thalamic circuits (Golding et al., 2014), such that these effects may be occurring at the post-transcriptional level or rely on extranuclear transcription, since they are not detected here.

Our results strongly suggest that subsets of HO neurons are developmentally co-opted from FO neurons in an input-dependent manner; this is also suggested by previous work in which FO neurons do not acquire FO identity and instead have a sustained HO identity. In principle, FO neurons could represent an entirely distinct population of neuron that mature late, but there is no evidence in birthdating studies for this: FO and HO nuclei are born simultaneously (see https://neurobirth.org/) (Baumann et al., 2023). Similarly, in principle, FO neurons could be born at the same time but migrate into the thalamus – the structure we are collecting – at later stages, giving a sampling-related illusion of late differentiation. Here, again, there is no evidence of such a migration occurring. Instead, the most parsimonious explanation is that FO neurons emerge from subsets of HO neurons to acquire new properties in an input-dependent manner. Noteworthily, HO neurons are still detected after the emergence of FO neurons, the latter being a transient event, suggesting that only subsets of HO neurons become FO neurons.

From an evolutionary standpoint, our findings thus support the idea that FO-type neurons could have developed from subsets of ancestral HO-type neurons. FO neurons might have been evolutionarily co-opted from some HO neuron types based on higher ability to transmit signals from high-resolution body receptors, perhaps reflecting their distinct metabolic, electrophysiological, and connectivity characteristics (Wong-Riley, 1979; Wong-Riley and Welt, 1980; Yamashita et al., 2013). It will be interesting in future studies to assess precisely how input-dependent processes regulate the gene regulatory networks at play in these cell-type-specific fate decisions.

## Materials and methods

### Utilization of Generative Al and Al-Assisted Technologies in the Writing Process

In the course of preparing this work, the authors employed Chat-GPT to enhance certain sections of the text. Following the utilization of this tool, the authors diligently reviewed and edited the content as required.

### Mouse Strains

The experiments were carried out in compliance with Swiss laws and obtained approval from both the Geneva Cantonal Veterinary Authorities and their ethics committee. The study strictly adhered to the ARRIVE guidelines. CD1 male and female mice were sourced from Charles River Laboratory. For embryonic experiments, mice were mated within an early morning 3-hour window at the Charles River facilities. In-house mattings were conducted overnight for post-natal experiments, with the following morning designated as time E0.5. Prior to the experiment, all mice were housed at the institutional animal facilities under animal caretaker control for their veterinary needs, covering the alternation of light and dark cycles, a 22°C temperature, and unlimited water and food provided. Transgenic mice consist of GAD67^GFP^ knock-in mice (Golding et al., 2014; Tamamaki et al., 2003).

### Surgical Procedures

#### FlashTag *in-utero* injection

FlashTag in utero injections were carried out as previously described (Govindan et al., 2018). Pregnant mice were anesthetized by isoflurane at gestational time point E11, positioned on a 37°C heated surgical platform for the duration of the surgeries. During surgeries, small incisions were made in the abdominal region to reveal the uterine horns, and one microliter of 10NM FlashTag (carboxyfluorescein succinimidyl ester, CellTrace CFSE, Life Technologies, C34554) was intracerebroventricularly injected by a nanoinjector equipped with pulled, beveled, and dust-cleaned glass pipet. Following the procedure, the uterine horns were carefully repositioned within the abdominal cavity, and sutures were used to close the peritoneum and skin. The mice were then placed on a warming pad until full recovery from anesthesia.

#### Bilateral enucleation

Bilateral enucleation surgeries were carried out on P0 pups as previously described (Golding et al., 2014). A small incision was made in the eyelid with a scalpel and the eye was separated from the optic nerve with scissors and removed from the orbit with forceps. Pups were placed in a box on a heating pad for recovery before returning to their mothers.

#### scRNA-seq collection and sequencing

Single-cell library captures were performed on between six or twelve pooled embryos, depending on the time point, with one library for each condition. The conditions are composed of dissociated cells and nuclei from the thalamic space of E11.5, E12.5, E13.5, E14.5, E16.5, E18.5 for the embryonic experiments (Fig. 1C), and independent dissections of LG, LP, VB, Po in both control and enucleation conditions for P3 and P7 time points (Fig. 5B).

#### Cell dissociation and FAC-sorting

Diencephalon of the embryos were collected in ice-cold Hank’s Balanced Salt Solution (HBSS) and segments were microdissected under a stereomicroscope. Thalamic chunks were then incubated in 200 *μ*l single-cell dissociation solution consisting of papain (1 mg/ml)-enriched HBSS at 37°C for 10 min with trituration every 2 min. Cells were further dissociated via gentle up-and-down pipetting and the reaction was stopped with the addition of 400 *μ*l of 2 mg/ml ovalbumin-enriched cold FACS buffer (2mg/mL glucose, 0.1% BSA, 1:50 Citrate Phosphate Dextrose from Sigma (C7165), 10U/mL DNase I, and 1μM MgCl2) and the cell suspension was then passed through a 40-*μ*m cell strainer to remove cellular aggregates. Cells were then centrifuged for 5 min at 500 ***g*** at 4°C. After removal of the liquid, the pellet was suspended in 250 *μ*l of cold FACS buffer and the resulting solution was finally sorted on a MoFloAstrios device (Beckman) to reach a concentration of 410 cells/*μ*l.

#### Nuclei isolation

Diencephalon of the embryos were collected in ice-cold Hank’s Balanced Salt Solution (HBSS) and segments were microdissected under a stereomicroscope. Thalamic chunks were frozen and stored at −80°C until further processing.

Nuclei were isolated using EZ Nuclei isolation kit as previously described (Santinha et al., 2023). In brief, the tissue was resuspended in 2 mL of ice-cold extraction buffer (Nuclei EZ prep, Sigma NUC101), gently dounce-homogenized (KIMBLE Dounce tissue grinder, Sigma D8938, 6 strokes with A and B pestle), and incubated for 5 minutes in a total of 4 ml EZ buffer.

The extracted nuclei were collected by centrifugation (500 x g at 4°C, 5 minutes) and supernatant removal followed by resuspension and incubation in EZ buffer for 5 min. After centrifugation, nuclear pellet was washed in 4 ml washing buffer containing 1% BSA in PBS, with 50U/ml of SUPERase-In (Thermo Fisher, cat# AM2696) and 50U/ml of RNasin (Promega, cat# N2611), centrifuged and nuclei were resuspended in 1 ml of washing buffer, filtered through a 30 μm strainer, stained with Hoechst (Invitrogen H3570, 1:500) for 5-10 minutes and processed for fluorescence-activated nuclei sorting on Beckman Coulter MoFlo Astrios. 20k nuclei were sorted and 42 *μ*l of nuclei suspension was used to load the 10x Genomics snRNA-seq preparation.

#### 10X Single cell RNA sequencing

Cell suspensions were captured using 10X Genomics Chromium Single Cell 3’ v3 reagents following the manufacturer’s protocol. The quality control of the cDNA and libraries was performed using Agilent’s 2100 Bioanalyzer. Subsequently, the libraries were sequenced using the HiSeq 4000 sequencer. The resulting FASTQ files obtained from the sequencing were processed and mapped with 10X Genomics Cell Ranger pipeline using the GRCm38 mouse genome as a reference. Default parameters for read mapping, counting and quality controls were used as described in the Cell Ranger V6.0.0 documentation.

#### C1 Single cell RNA sequencing

Cells stained with FlashTag were captured using the C1-Fluidigm system. Following the previously mentioned dissociation protocol, 8 *μ*L of cell suspension was sorted on a MoFloAstrios to enrich for the top 20% most FlashTag-positive cells. Each sample representing a different embryonic time points was mixed with C1 suspension reagents (2 *μ*L; Fluidigm) for a total cell suspension volume of approximately 10 *μ*L, containing ∼500 cells per *μ*L. This mixture was then loaded onto the C1 Single-Cell AutoPrep integrated fluidic circuit (HT-800, Fluidigm, 100-57-80). cDNA synthesis and preamplifications were conducted following the manufacturer’s instructions for the C1 system (Fluidigm). Captured cells were imaged using the ImageXpress Micro Widefield High Content Screening System (Molecular Devices). Single-cell RNA-sequencing libraries of the cDNA were prepared using the Nextera XT DNA library prep kit (Illumina). Libraries were multiplexed and sequenced in accordance with the manufacturer’s recommendations, employing paired-end reads on the HiSeq4000 platform (Illumina).

#### Single cell RNA-seq Data Analysis

All single cell transcriptomics analyses were performed using R Statistical Software (v4.2.2), the Seurat V4 package and Bioconductor packages (Hao et al., 2021; Huber et al., 2015; R: The R Project for Statistical Computing). Graphs and visualization were generated with the ggplot2 package (Wickham).

For quality control, independent libraries were processed as Seurat objects. All cells with fewer than 200 detected genes and genes expressed in fewer than 3 cells were removed. In detail, we were able to retrieve 614 FlashTag cells at E11.5, 310 at E12.5, 303 at E13.5, 241 at E14.5, 93 at E18.5, for the FlashTag condition, and 2066 at E11.5, 2449 at E12.5, 2658 at E13.5, 3013 at E14.5, 3652 at E16.5, and 3578 at E18.5 for non-FlashTag nuclei (Fig. 1C). PCA was performed on variable genes to reduce dimensionality of the dataset. UMAP was based on the reduced dimensional space of the 20 most significant dimensions of the PCA using the UMAP package Barnes-Hut version of Seurat with a perplexity set at 30 (Fig1. C). A UMAP-based clustering analysis was then performed by the shared nearest neighbor (SNN) modularity optimization algorithm (van der Maaten, 2014). The number of independent genes detected ranged between 2684 and 3994 with an average of 3148 by cluster. Differentially expressed genes between clusters were obtained by Seurat-implemented Wilcoxon rank sum tests with default parameters. The cluster identities in this UMAP-space were uncovered by feature plots of typical cell type marker genes (Fig. 1E,F). The first three embedded dimensions of the UMAP were output for further use to represent the pattern of various features during differentiation. The minimal distance parameter of the UMAP was set to 0.3 and the number of neighboring points used in local approximations was set to 30.

#### Merging of dataset

Individual datasets were integrated together following the scRNA-seq integration vignette of Seurat documentation with following steps. Individual experiments were normalized by the NormalizeData Seurat function with default parameters, and 2000 variable features were computed with the “vst” selection method. Integration anchors were then computed for 30 dimensions with a k-filter parameter of 50, and used for the final integration.

#### Pseudotime projection

Pseudodifferentation and pseudotime scores were assigned using the monocle 3 R library following the published pipeline, with default parameters (Fig. 2A).

Pseudo-time analysis was performed with Monocle 3 using genes that have passed the quality control of the Seurat object creation (Qiu et al., 2017a). Genes considered as defining the progression of the pseudo-time were those that were detected as having an expression above 0.5 by Monocle 3. Negative binomial was considered for the model encoding the distribution that describes all genes. During the pseudo-time processing, the dimensionality of the dataset was reduced by the Discriminative Dimensionality Reduction with Trees (DDRTree) algorithm on the log-normalized dataset with ten dimensions considered.

#### Transcriptional waves

Transcriptional wave analysis was performed with default parameters as described previously (Telley et al., 2016) (Fig. 2B). Genes presenting interesting variations were regrouped along a pseudo-time axis, forming clusters composed of similar time-dependent gene expressions. These clusters of patterns were labeled as waves and processed for further GO term enrichment (Fig. 2C).

GO term, pathways enrichment analysis, and their representations were performed using the R pathfindR package (Ulgen et al., 2019), in accordance with the library documentation and DAVID bioinformatics resources 6.8. (Huang et al., 2008, 2009), on list of differentially expressed genes obtained by *findmarker* and *findallmarker* Seurat functions (Fig. 1F and 2C). Gene ontology networks are a follow-up analysis of the Seurat lists of differentially expressed genes. After the list creation by the *findmarker* Seurat function with default parameters, the gene names, log-fold changes, and adjusted p-values are inputted into the run_pathfindR function of the pathfinder R package (v 2.2.0), with the following parameters: custom_genes defined as mmu_kegg_genes and custom_descriptions = mmu_kegg_descriptions, as described in the mice-specific PathfindR workflow (https://cran.r-project.org/web/packages/pathfindR/vignettes/non_hs_analysis.html). Then, the combine_pathfindR_results function was used to compare results of different conditions and obtain specific and common pathways (Fig. S5B; S6A,B; S7B,C,D).

#### Spatial Transcriptomics and Label Transfer

A coronal representation of a whole-brain spatial transcriptomics experiment dataset was obtained (Zhang et al., 2023) and imported into R. Coronal level 37 was selected as it is representative of the studied thalamic space (Fig. S9D). The LG P7 control and deprivation clusters, as previously described (Fig. 5B), were integrated following the merging of datasets procedure outlined in the previous paragraph. The control condition served as the reference, and its subclustering labels (Seurat clusters 0 to 7) were transferred to the deprivation dataset (Fig. S9A). Genes markers of these control sub-clusters that were common to expressed genes in the spatial dataset (all genes specific to each sub-cluster with a positive log fold-change after the *findallmarker* Seurat function) were used to compute a pseudogene for each subcluster based on their mean expression. These pseudogenes were then represented for both the control and deprivation conditions (Fig. S9C, left part). Subsequently, the representative expressions of these sub-clusters were individually plotted along the pseudotime values of each cell (Fig. S9C, middle) and on the spatial dataset (Fig. S9C, right).

#### In situ hybridizations

High resolution in situ hybridization images were selected and extracted from the Allen Brain Atlas using R scripts to connect and download from the Allen Brain Atlas Application Programming Interface (http://mouse.brain-map.org/static/api) (Fig. 1G; 2D).

#### Detection of embryonic LG, LP, VB and PO cells based on mature markers

Initially, the top expressed genes for FO, HO, LG, VB, Po, LP were identified through literature references (Frangeul et al., 2016) (Table S6; Table S10). Pseudogene expression was subsequently computed for each modality based on these markers. For terminal E18 clusters, the most promising cells in each modality were selected by considering the higher expression quantile within their respective categories, ultimately representing core identity cells (Fig. 3A). Specific markers for these candidate cells were computed, with the top 30 markers selected in a second pass. To address expression level imbalances between modality pairs, a normalization of the 2nd order pseudogene expression was performed, scaling values between 0 and 10. Ratios of normalized values for modality pairs were then calculated. The quantile selection of the best cells was repeated, resulting in the final selection of cells representing their respective modalities (Fig. 3C; 4C,D). Finally, an unsupervised clustering analysis was conducted in the embryonic UMAP space, revealing distinct cell groups with specific identities. In this UMAP space, pseudogene expression based on the top markers was computed for each modality.

## Acknowledgements

We thank the Genomics Platform and FACS Facility of the University of Geneva; A. Benoit for technical assistance; all members of the Jabaudon laboratory for constructive exchanges during the project.

## Author Contributions

Conceptualization: D.J.; Methodology: Q.L.G., R.W., P.A., L.F.; Formal analysis: Q.L.G.; Writing – original draft: D.J.; Writing – review & editing: Q.L.G., L.F., D.J.; Visualization: Q.L.G., L.F., D.J.; Supervision: D.J.; Project administration: D.J.; Funding acquisition: D.J.

## Competing Interests

The authors declare no competing or financial interests.

## Funding

The Jabaudon laboratory is supported by the Swiss National Science Foundation, the Carigest Foundation, the Société Académique de Genève FOREMANE Fund, and the European Research Council. P.A. was supported by EMBO (ALTF 136-2017) and IRP (P 185F) Postdoc Fellowship. R.J.W. was supported by the Deutsche Forschungsgemeinschaft (DFG) Wa 3783/1.

## Data Availability

Single-cell RNA sequencing data generated in this study has been deposited in NCBI Gene Expression Omnibus (GEO) under accession number XXX. This paper does not include proprietary code.

## Supplementary Figures

**Fig. S1.**
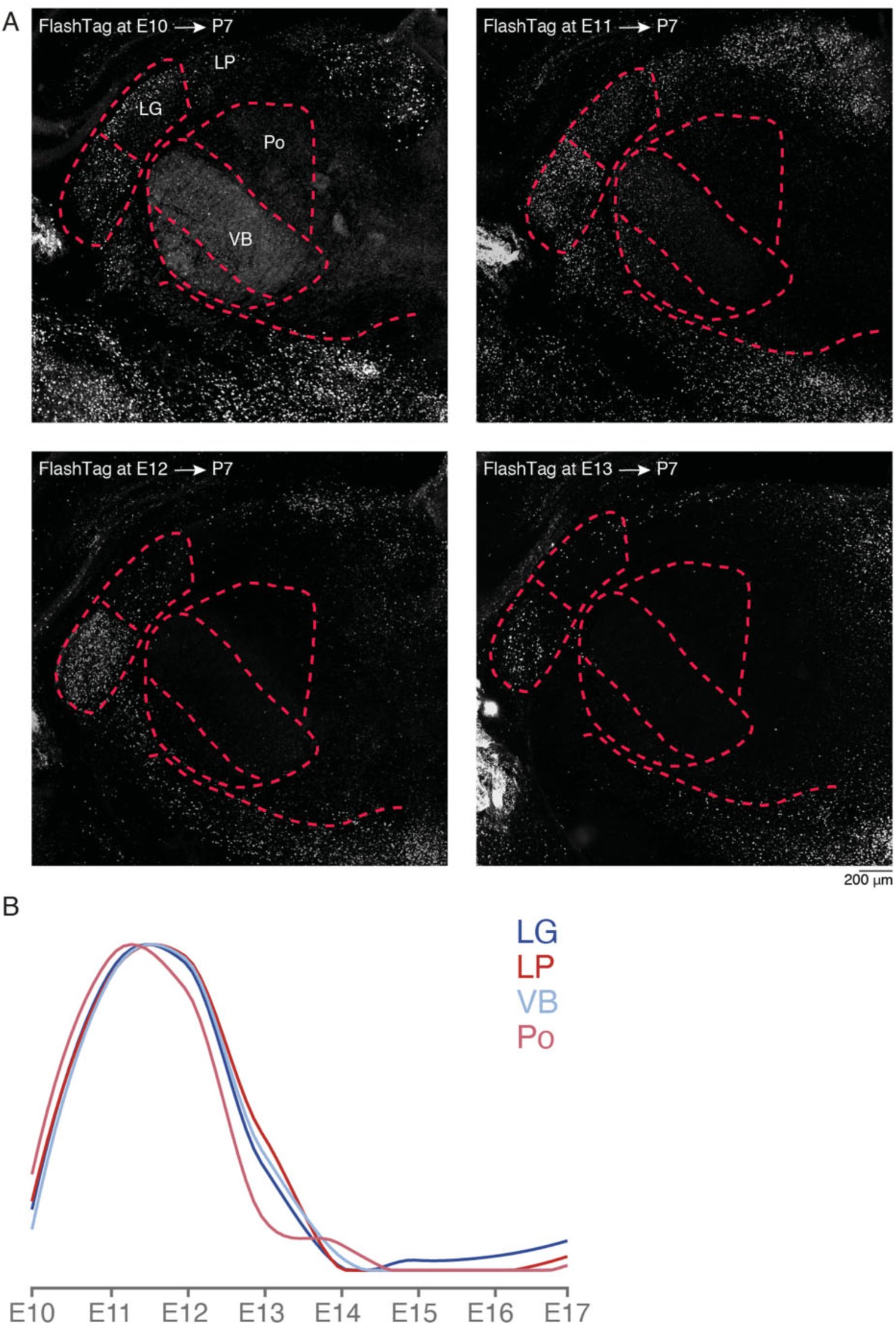
Thalamic identity of VZ-born neurons between E10 and E13. (A) Example images showing thalamic nuclei of P7 pups injected with FlashTag at different embryonic stages. (B) Line plot displaying the birth date of VB, Po, LG and LP thalamic nuclei (data from https://neurobirth.org/) (Baumann et al., 2023). LG, dorsolateral geniculate nucleus; LP, pulvinar/latero-posterior nucleus; Po, posteromedial nucleus; VB, ventrobasal nucleus. Scale bar: 200 μm (A).

**Fig. S2.**
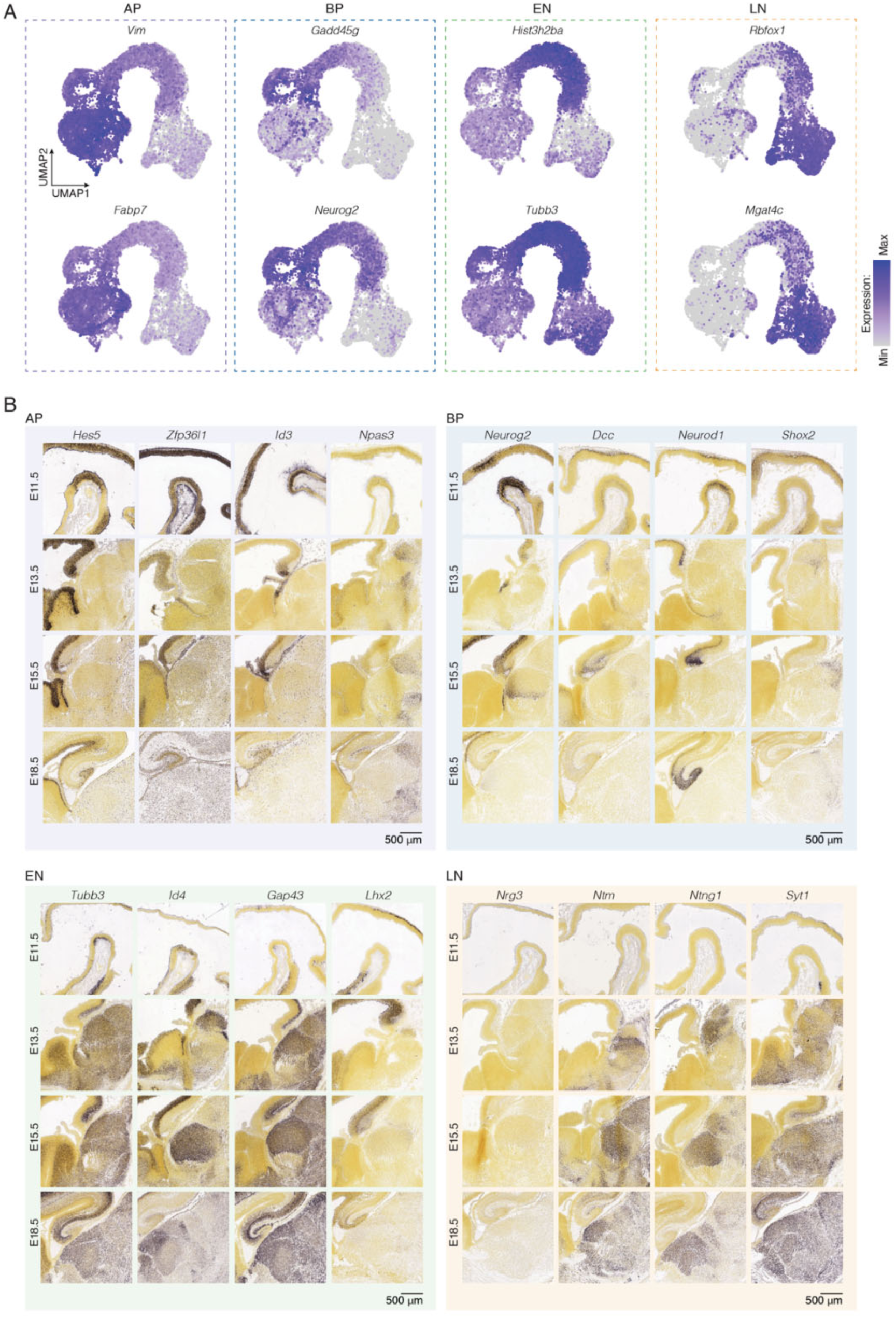
Expression of select genes in AP, BP, EN and LN. (A) Example feature plot of select genes for apical progenitors (AP), basal progenitors (BP), early neurons (EN) and late neurons (LN). (B) In situ hybridization (ISH) sections of selected differentially expressed genes showing distinctive expression during mouse embryonic development; image source: Allen Developing Mouse Brain Atlas (developingmouse.brain-map.org). Scale bar: 500 μm (B).

**Fig. S3.**
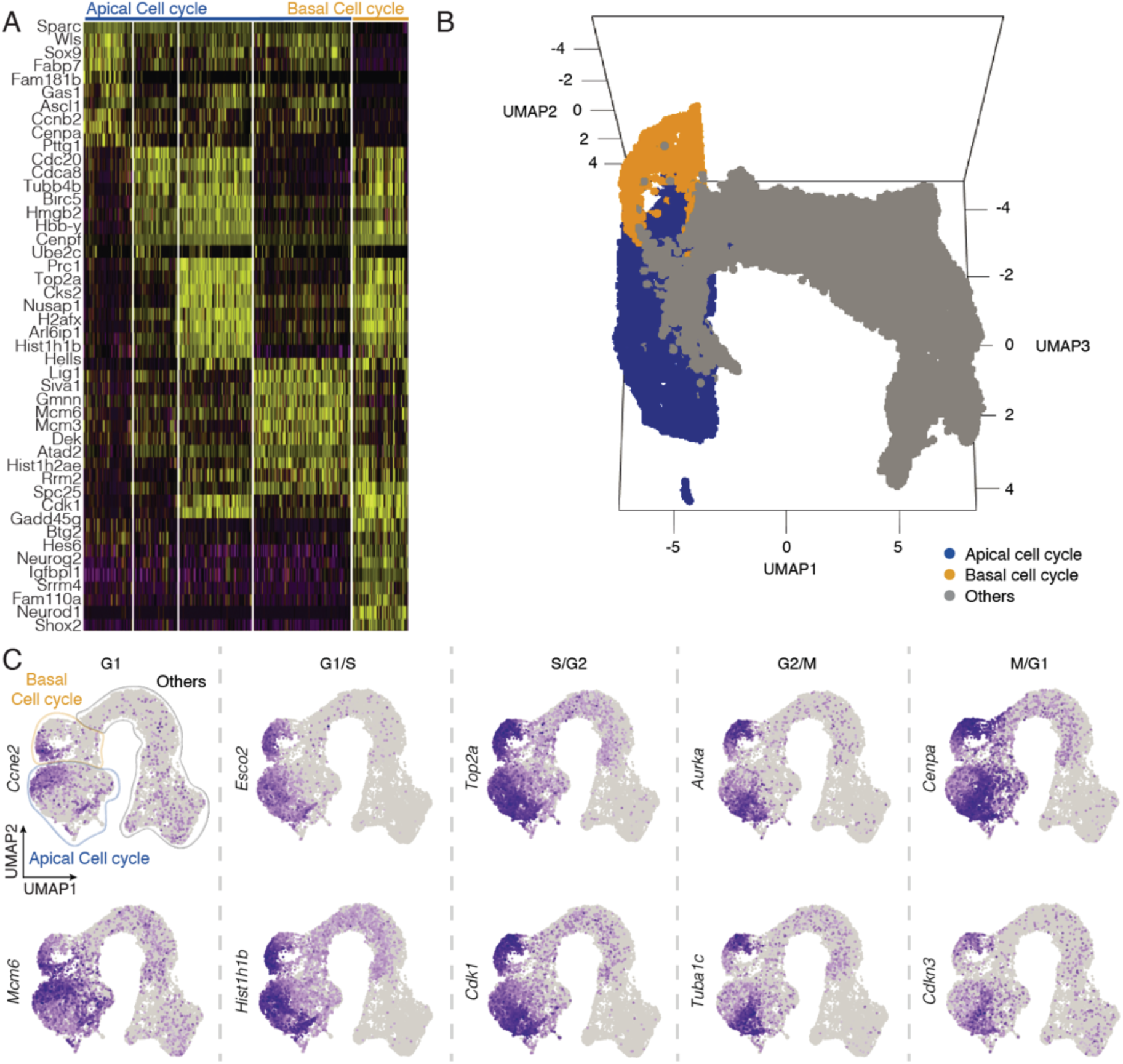
Molecular signature of apical and basal cell cycle in the developing thalamus. (A) Expression of the top 10 of each cell cycle phase highlight the presence of apical and basal cell cycle. (B) UMAP representation in 3D showing the apical and basal cell cycle. (C) Example feature plot of selected genes for each cell cycle phase.

**Fig. S4.**
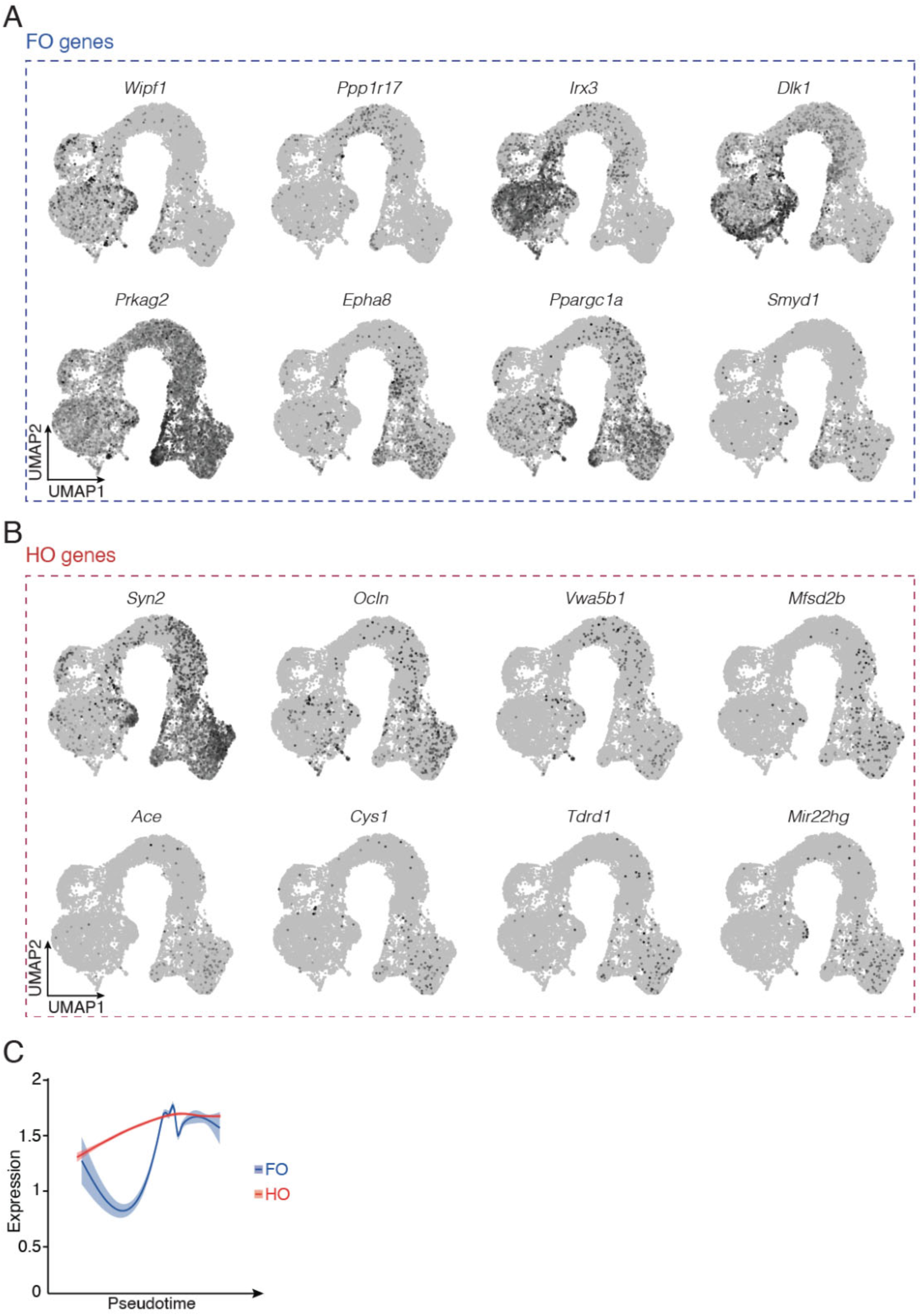
Expression of select genes in FO and HO thalamic neurons. (A) Example feature plot of select genes for FO genes. (B) Example feature plot of select genes for HO genes. (C) Line plot showing the dynamics of average expression of shared genes along pseudotime between FO and HO. FO, first order; HO, higher order.

**Fig. S5.**
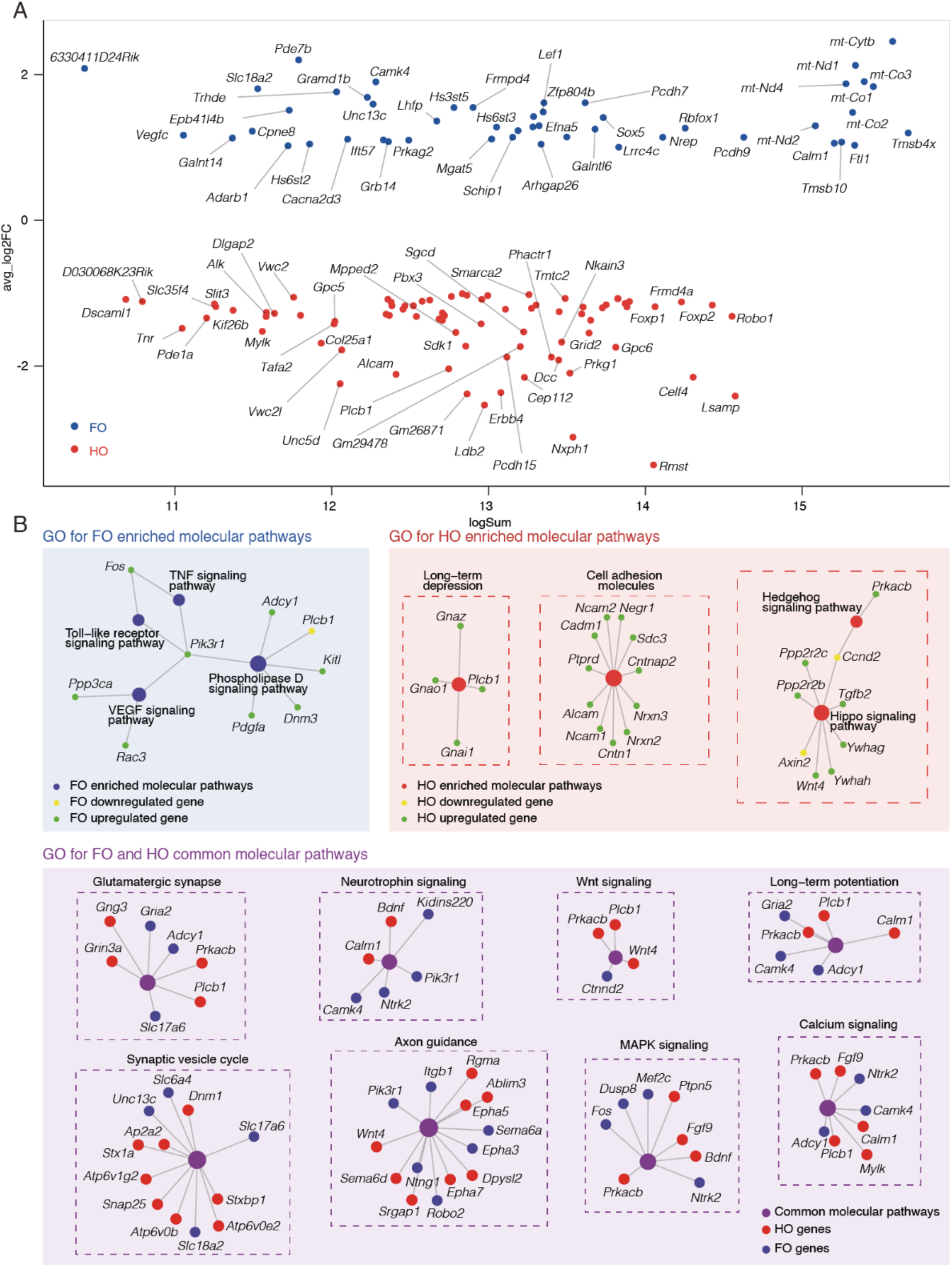
Molecular signature and gene ontology of FO and HO thalamic neurons. (A) MA plot showing log fold-change (y-axis) compared with mean expression value (x-axis) (log Fold-Change >1 for FO, <(−1) for HO, logSum > 8 and p-value < 0.01), negative fold change representing specific genes in HO (red) and positive fold change representing specific genes in FO (blue). (B) String plots representing gene ontology networks for FO enriched molecular pathways (blue), HO enriched molecular pathways (red) and FO and HO common molecular pathways (purple). FO, first order; HO, higher order.

**Fig. S6.**
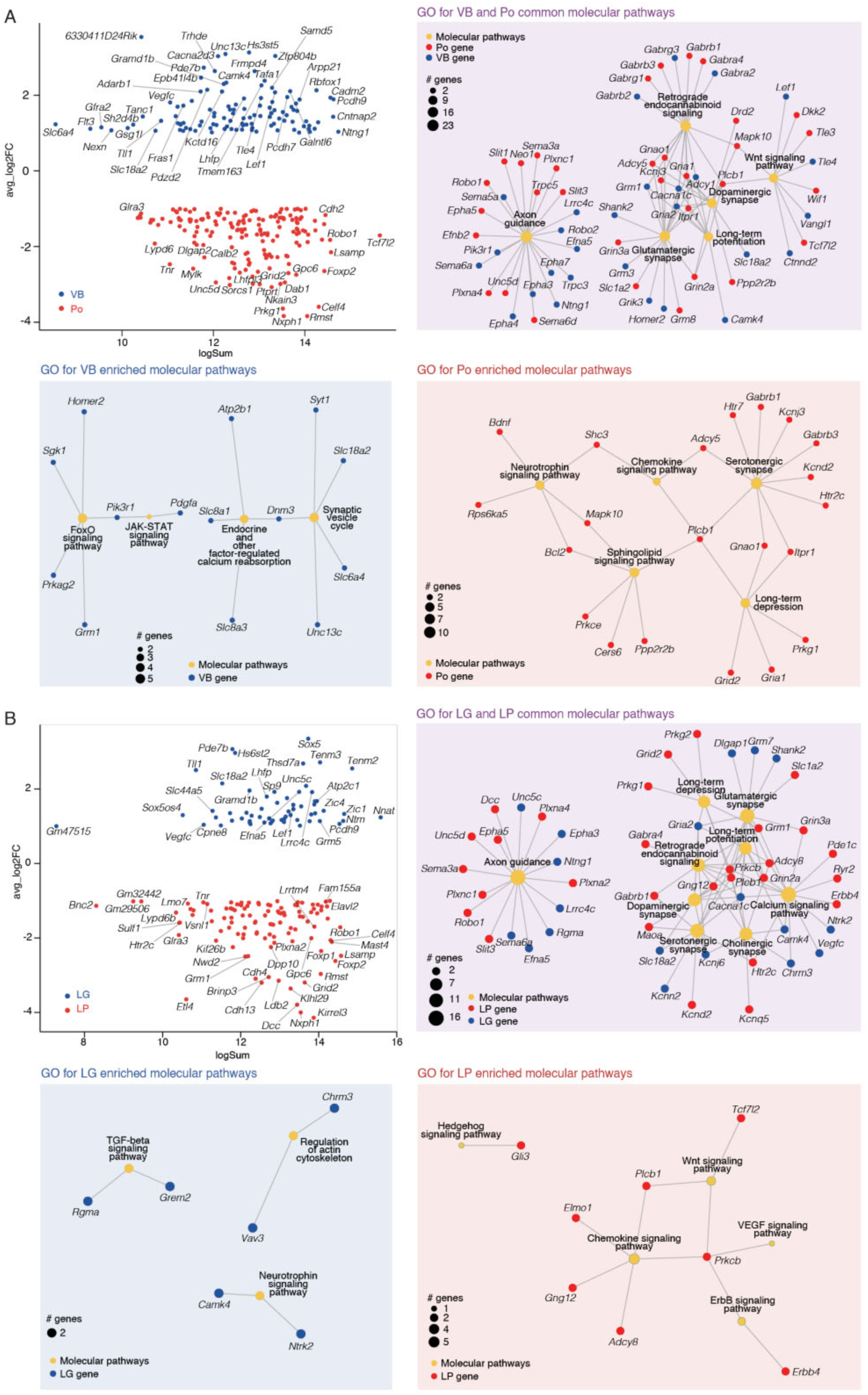
Molecular signature and gene ontology of nucleus-specific thalamic neurons. (A) MA plot showing log fold-change (y-axis) compared with mean expression value (x-axis) (same criteria as in Fig. S5), negative fold change representing specific genes in Po (red) and positive fold change representing specific genes in VB (blue). String plots representing gene ontology networks for VB enriched molecular pathways (blue), Po enriched molecular pathways (red) and VB and Po common molecular pathways (purple). (B) MA plot showing log fold-change (y-axis) compared with mean expression value (x-axis), negative fold change representing specific genes in LP (red) and positive fold change representing specific genes in LG (blue). String plots representing gene ontology networks for LG enriched molecular pathways (blue), LP enriched molecular pathways (red) and LG and LP common molecular pathways (purple). LG, dorsolateral geniculate nucleus; LP, pulvinar/latero-posterior nucleus; Po, posteromedial nucleus; VB, ventrobasal nucleus.

**Fig. S7.**
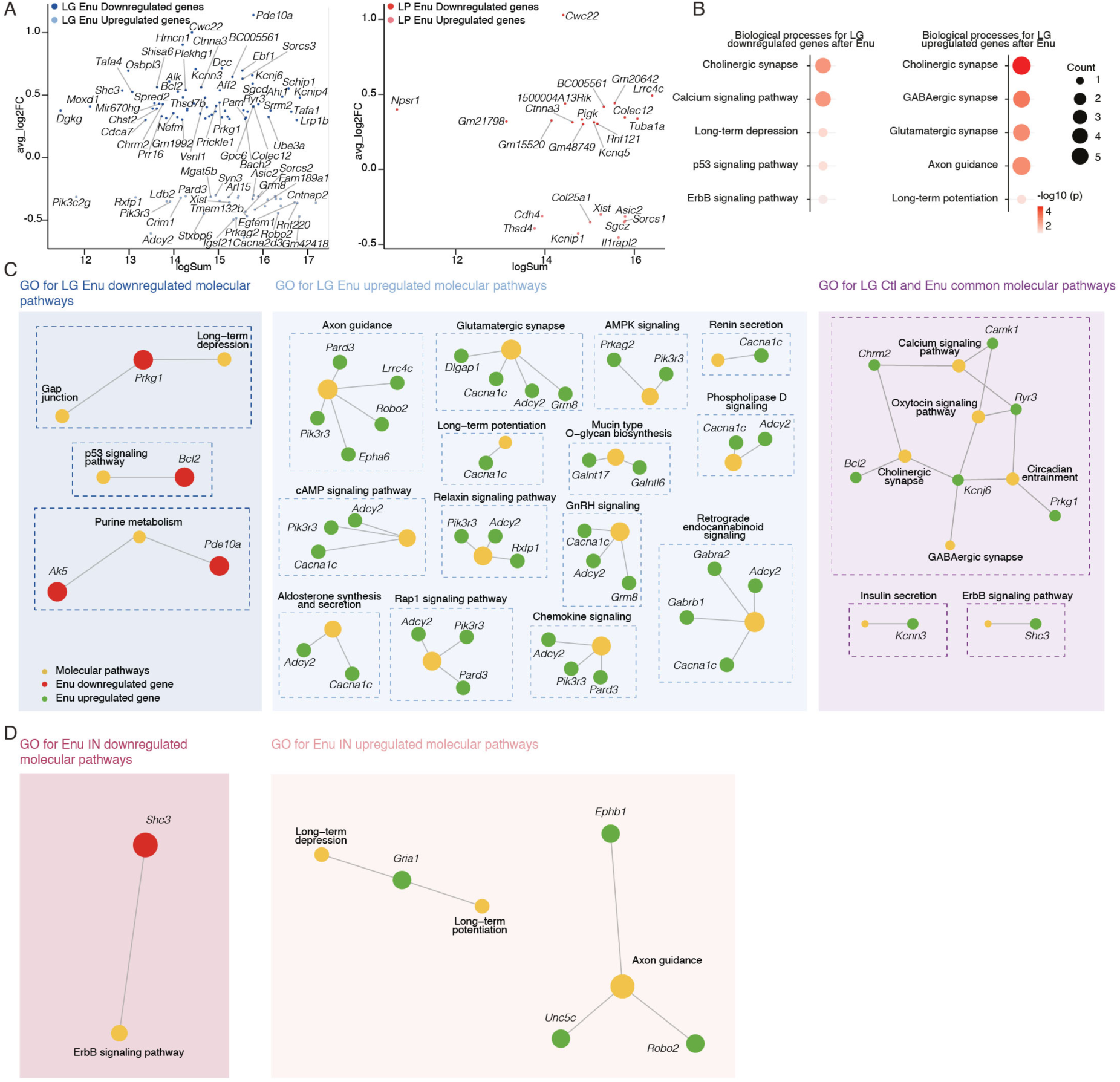
Molecular signature and gene ontology of modified by input deprivation. (A) MA plot showing log fold-change (y-axis) compared with mean expression value (x-axis) (same filtering criteria as Fig. S5), negative fold change representing up-regulated genes in LG Enu (light blue) and positive fold change representing down-regulated genes in LG Enu (dark blue). (B) Example of biological processes of gene ontologies associated with LG downregulated genes after enucleation (left) and LG upregulated genes after enucleation (right). (C) String plots representing gene ontology networks for LG Enu downregulated molecular pathways (dark blue), LG Enu upregulated molecular pathways (light blue) and LG Ctl and Enu common molecular pathways (purple). (D) String plots showing gene ontology networks for IN downregulated molecular pathways (dark red), IN Enu upregulated molecular pathways (light red). Ctl, control; Enu, Enucleation; IN, interneuron; LG, dorsolateral geniculate nucleus; LP, pulvinar/latero-posterior nucleus.

**Fig. S8.**
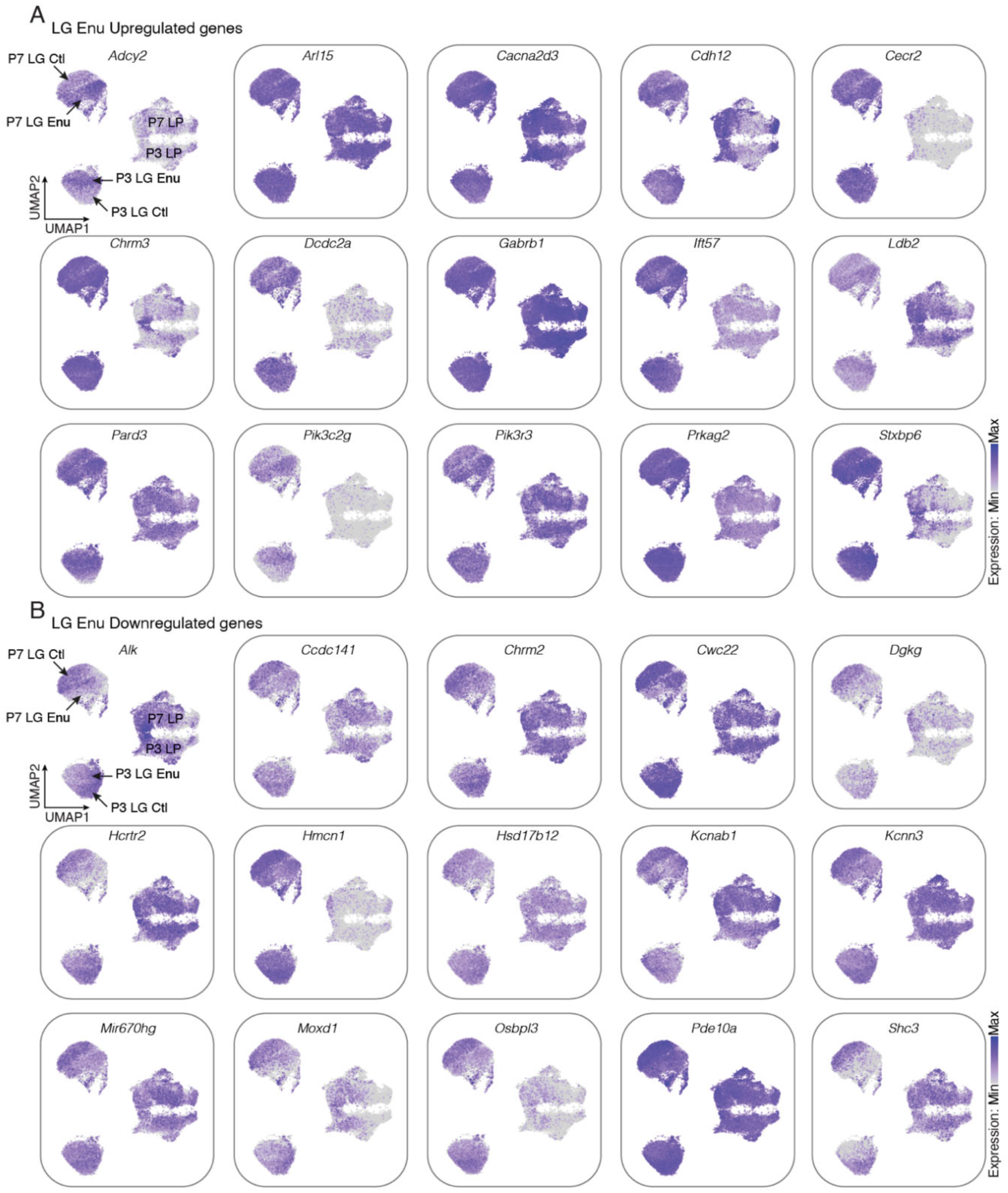
Expression of select genes modified by input deprivation. (A) Example feature plot of select genes upregulated in LG after enucleation. (B) Example feature plot of select genes downregulated in LG after enucleation. Ctl, control; Enu, Enucleation; LG, dorsolateral geniculate nucleus.

**Fig. S9.**
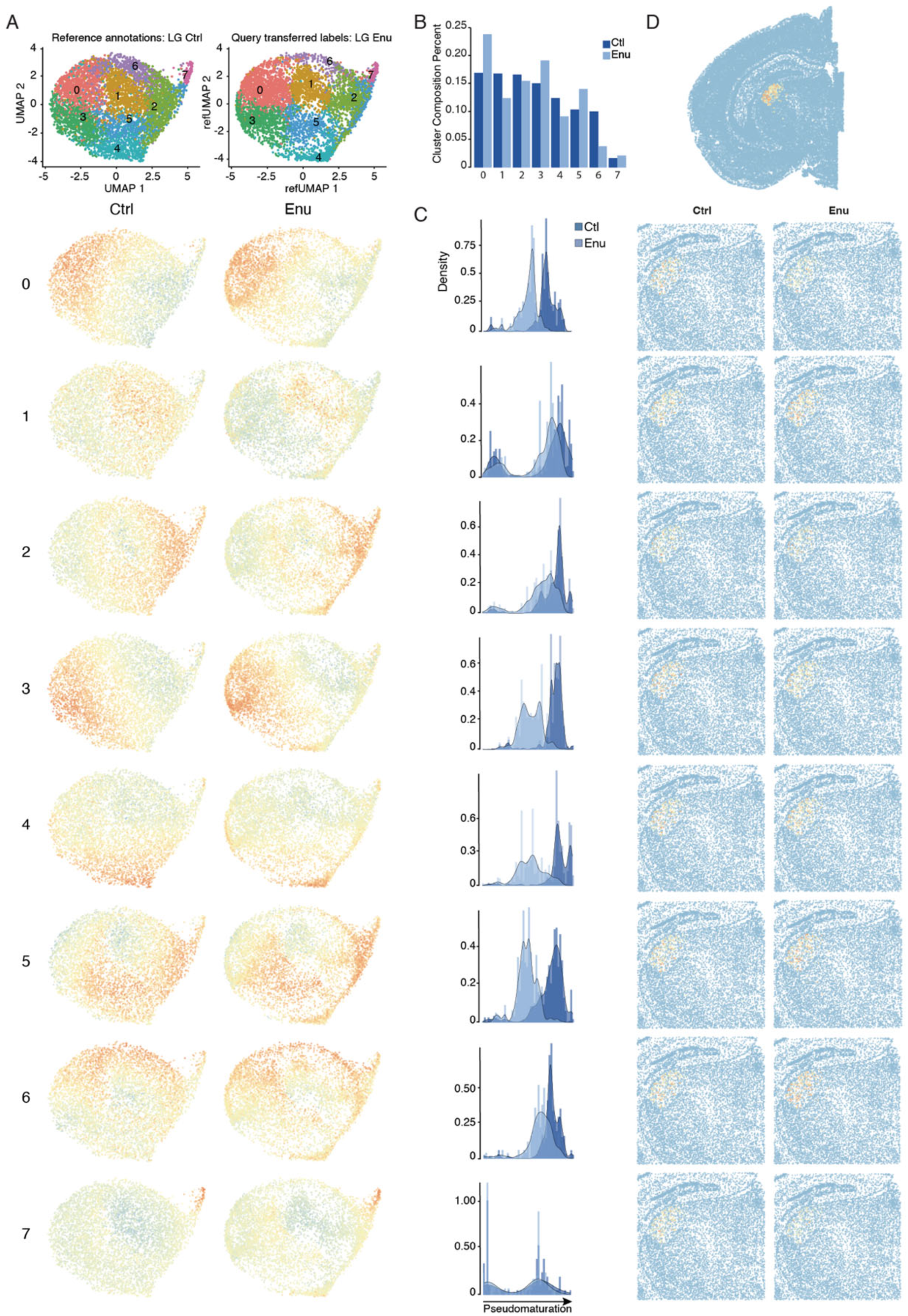
Cell-type-specific transcriptional responses to input deprivation within LG. (A) UMAP representation of the integration of P7 LG Enu on the P7 LG Ctl embedding. unbiased clustering of the integrated deprivation and control P7 LG space (cf methods). (B) Representation of cluster composition in Ctl and Enu condition displayed in a histogram. (C) UMAP representation of either LG Ctl or LG Enu condition for each identified cluster along the pseudotime axis. (D) Spatial transcriptomic of differentially expressed genes of LG subclusters (spatial transcriptomic dataset from Zhang et al., 2023). Ctl, control; Enu, Enucleation; LG, dorsolateral geniculate nucleus.

## Supplementary Tables

**Table S1. Apical progenitors, basal progenitors, early neurons and late neurons differentially expressed genes list (related to Fig. 1E).**

**Table S2. Apical and basal cell cycle progenitors differentially expressed genes list (related to Fig. S3).**

**Table S3. Apical progenitors, basal progenitors, early neurons and late neurons gene ontology analysis (related to Fig. 1F).**

**Table S4. Gene expression across transcriptional waves along pseudotime (related to Fig. 2B).**

**Table S5. Gene ontologies associated with transcriptional waves (related to Fig. 2C).**

**Table S6. List of mature FO and HO genes used to define new differentially expressed genes (related to Fig. 3B).**

**Table S7. FO and HO new differentially expressed genes list (related to Fig. 3B).**

**Table S8. FO and HO waves gene list (related to Fig. 3D).**

**Table S9. Biological processes associated with FO and HO transcriptional maps clusters (related to Fig. S5).**

**Table S10. List of mature VB, Po, LG and LP genes used to define new differentially expressed genes (related to Fig. 4A and B).**

**Table S11. VB, Po, LG and LP new differentially expressed genes list (related to Fig. 4A, B and G).**

**Table S12. VB, Po, LG and LP waves gene list (related to Fig. 4E and F).**

**Table S13. Biological processes associated with VB, Po, LG and LP transcriptional maps clusters (related to Fig. S6).**

**Table S14. LG and LP Ctl vs. Enu differentially expressed genes list (related to Fig. 5B, S7 and S8).**

**Table S15. Biological processes associated with LG Ctl, LG Enu, LP Ctl and LP Enu transcriptional maps clusters (related to Fig. S7).**

**Table S16. LG subclusters differentially expressed genes list (related to Fig. S9).**

## Notes

### Competing Interest Statement

The authors have declared no competing interest.

